# Structuring effects of archaeal replication origins

**DOI:** 10.1101/2023.11.15.567178

**Authors:** Clémence Mottez, Romain Puech, Didier Flament, Hannu Myllykallio

## Abstract

Archaea use eukaryotic-like DNA replication proteins to duplicate circular chromosomes similar to those of bacteria. Although archaeal replication origins have been maintained during the evolution, they are non-essential under laboratory conditions. Here we propose the local deviations from Chargaff’s second parity rule of archaeal chromosomes result from the biased gene orientation and not from mutational biases. Our computational and experimental analyses indicate that the archaeal replication origins prevent head-to-head collisions of replication and transcription complexes as well as participate in coordination of the transfer of genetic information. Our results therefore suggest that the archaeal replication origins have alternative functions not related to their role in initiation of DNA replication.

---

Archaea are a fascinating group of micro-organisms with considerable evolutionary, environmental, and biotechnological interest since the pioneering work of Woese and Fox in the 1970s. From a molecular mechanistic point of view, studies on archaeal DNA replication have attracted extensive interest since the publication of the first archaeal genome sequence, which revealed that the architecture of archaeal circular chromosomes is very similar to that of bacteria in terms of gene density and operon structures. Nevertheless, archaeal replication proteins are more closely related to their eukaryotic, and not bacterial, counterparts^1^. Strikingly, many archaeal DNA replication proteins including DNA primase, replicative helicase, and DNA polymerase are evolutionary unrelated in bacteria and archaea raising questions about how functional parallels between semi-conservative and bidirectional DNA have evolved in two prokaryotic domains^2^.

Surprisingly, despite the overall structure of archaeal replication origins has been maintained during the evolution^3^, these archaeal sequence elements required for the site-specific initiation of DNA replication are non-essential^4, 5^. This raises a possibility that archaeal replication origins have alternative functions not related to replication initiation but that are potentially required for long-term viability under natural conditions.

These observations prompted us to obtain quantitative data for the genome composition of completed archaeal genome sequences as a proxy to understand the diversity of the archaeal genome structure. Erwin Chargaff’s parity rule 1 states that, in double-stranded DNA, the molar ratios of guanine and cytosine as well as adenine and thymidine are identical, which simply reflects base pairing in the DNA duplex. Later, he extended this observation to his second parity rule, indicating that this holds even for individual strands of dsDNA genomes^6^. The basis for this conserved phenomenon, except for mitochondria and ssDNA viruses, remains poorly understood. It is even argued that this DNA sequence symmetry has no biological basis but arises from randomness^7^. However, at the whole genome level, local deviations in nucleotide composition result in asymmetries in base composition and locally violate the second parity rule. These are referred to as nucleotide “skews” indicating for instance the local excess of guanine over cytosine that can be presented as (G - C)/(G + C) in a given genome window. These local variations in the second parity rule have biological origins. In particular, combinations of strand-specific biases in DNA replication from single or multiple replication origins, transcription, gene density, and codon biases contribute to deviations from the second parity rule in^8^. It has also been demonstrated that strand-biased cytosine deamination causing cytosine-to-thymine mutations at the replication fork contributes to GC skew. Since the lagging strand of the replication is exposed in ssDNA form at the fork, the rate of cytosine deamination is increased. This has been experimentally demonstrated using accelerated laboratory evolution experiments using cytosine deaminase as a strand-specific DNA mutator^9^. The GC skew phenomenon can also be used to map the transition points between the leading and lagging strands that correspond to the replication origins and termini^10^.

We first confirmed a strong linear correlation (R^2^=0.9997) between cytosine and guanine counts for single strands of available archaeal replicons, with an approximate mean size of 2 Mb (**Fig. 1a**). This agrees well with the earlier analysis using 170 archaeal sequences (R^2^=0.99)^7^, further indicating that archaeal genomes follow Chargaff’s second parity rule. We next quantified the extent of the GC skew in archaeal genomes, which has not been systematically investigated previously. This is surprising considering that the first archaeal replication origins were predicted using GC skew more than 20 years ago^11^, followed by experimental confirmation^12^. To this end, we implemented the Skew Index Test^13^ (SkewIT) for archaeal genomes (for compilation of our results, see 10.5281/zenodo.8126182), which was originally used for large-scale analyses of bacterial genomes. This index provides a single numerical value [Skew Index (SkewI)], ranging from 0 to 1, which presents the degree of GC skewness of the complete archaeal genomes. **Fig. 1b** indicates that archaeal genomes indicate positive SkewI values with a mean value of 0.27±0.15 (S.D, n=807). However, this value is significantly lower than that previously reported for bacterial genomes (0.82±0.22, n=15067). We did not observe a major variation in the SkewI index between the different archaeal groups, which typically range between 0.21 and 0.28. This suggests that the different ploidy numbers of archaeal species^14^ do not modulate archaeal GC skew. Nevertheless, many archaeal genomes, corresponding to SkewI values higher than 0.5, were detected, particularly for euryarcheota, which presented the majority of the data points (n=543). Interestingly, cyanobacteria behaved in these analyses very similarly to archaeal species (0.24±0.22, n=153), but very different from the bacterial averages, as cyanobacterial SkewI was significantly lower (P-value < 0.001) than the other bacteria. Average values observed for *Synechocales* were skewed towards higher values and were, on average higher (0.36±0,2847, p-value < 0.002, n=64), indicating variability when compared with the other cyanobacterial phyla. The parallel between archaea and cyanobacteria is of interest, as DNA replication initiators or replication origins are not essential in either phylogenetic group^4, 15^, as, in both cases, the use of multiple replication origins and alternative replication mechanisms has been suggested. Notably, in archaea, recombination-associated DNA synthesis has been biochemically reconstituted using DNA polymerases (PolD and/or PolB) and the recombinase RadA suggesting that interplay between origin(s) and recombination-dependent mechanisms can be used to initiate DNA replication in archaea^16^.

**Figure 1.**
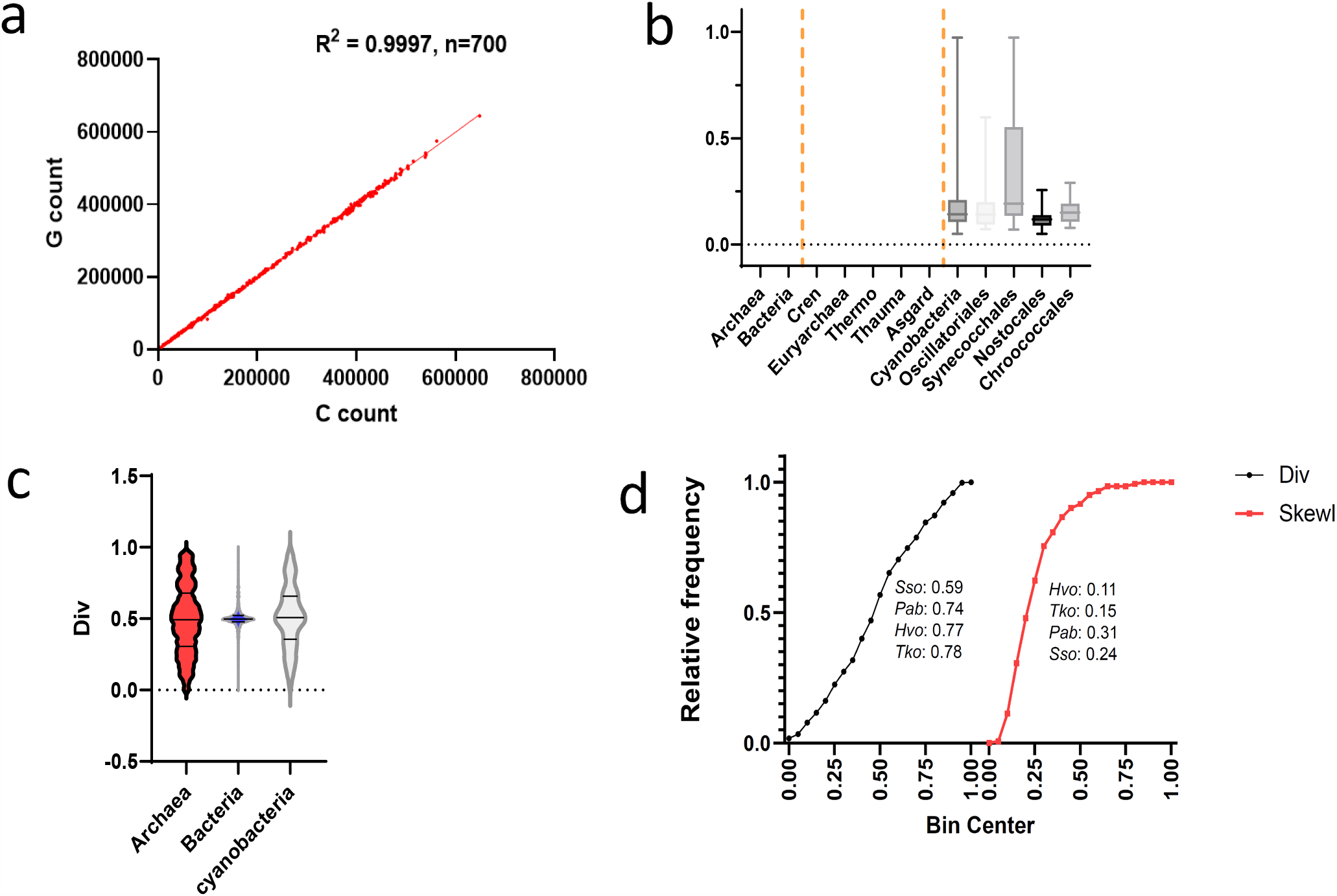
Archaeal genome analyses are based on local deviations from Chargaff’s second-parity rule. ***a)*** Counts of G vs. C on a single DNA strand for all archaeal genomes are plotted. This plot confirms that Chargaff parity rule 2 is true for archaea. ***b)*** Violin plot indicating SkewI values for all archaeal genomes using 15,067 bacterial genomes representing 4,471 species and 1,148 genera as comparison points. Unpaired two-tailed t-tests with Welch’s corrections were used to determine the statistical significance of the observed differences (see the text). ***c)*** Distribution of Div values corresponding to the fraction of the predicted leading strand from the fits of the GC skew for archaeal, bacterial, and cyanobacterial genomes. The mean values for the archaeal, bacterial, and cyanobacterial strains were 0.495, 0.501, and 0.511, respectively, which were not statistically different. However, Levene’s and Bartlett’s tests (XLSTAT 2023.1.6.1410) revealed that the variances between the archaeal and bacterial data sets were significantly different (P-value < 0.0001) ***d)*** Cumulative frequency distributions of Div and SkewIT values for archaeal genomes. Values for *Sulfolobus solfataricus* (*Sso*), *Pyrococcus abyssi* (*Pab*), *Haloferax volcanii* (*Hvo*), and *Thermococcus kodakerensis* (*Tko*) chromosomes are shown in the graph.

We further quantified the GC skew in the archaeal genomes (**Fig. 1c**). This has recently been facilitated by the establishment of the Skew Database (SkewDB), which includes precalculated GC skews for more than 30,000 bacterial and archaeal chromosomes and plasmids larger than 100 kb^13^. The obtained skews were calculated in windows of 4096 nucleotides and fitted using different mathematical models. These included the relative excess of G over on the predicted leading and lagging strands while the fraction of the chromosome replicated as the leading strand was denoted as “div.” For archaea, SkewDB indicates an average of 7-8 excess Gs in 1000 base windows, which is much lower than that observed for bacteria (23-25 excess G nucleotides)^17^. The fold change between bacterial and archaeal SkewI values was approximately 3.42, further indicating that the GC skew phenomenon is conserved in archaea, albeit to a lower extent than in bacteria. The lower amplitude of GC skew archaea agrees well with why the use of cumulative GC skew facilitates the prediction of archaeal replication origins. We also plotted the archaeal “div” values that reported the fraction of the predicted leading strand from the fits of the GC skew. For this plot, we observe that bacterial values (except for cyanobacteria) are clearly centered around the value of 0.5, which indicates that equally sized replicons initiate from a single well-defined replication origin and terminate at well-defined chromosomal sites. However, for archaeal and cyanobacterial chromosomes revealed a significant variance in “div” values between archaea/cyanobacteria and the other bacteria. Although the mean values for the different datasets were not significantly different, Bartlett’s and Levene’s tests revealed that the variances of the div-values between the bacteria and archaea groups were not homogenous (P-value < 0.0001). This observation is consistent with the use of multiple replication origins, alternative replication initiation mechanism and/or poorly defined replication termination zones in many archaea and cyanobacteria.

**Fig. 2a** depicts the replicon structure with cumulative GC and transcriptional strand bias skews for *Methanobrevibacter arboriphilis* strain SA as a representative of an “ideal” archaeal chromosome with SkewI and div values of 0.79 and 0.51, respectively. The observed skews and values indicate that in the archaeal species with bacterial-like GC-skew, replication, and transcription are typically organized in the same direction to minimize genotoxic head-on collisions of the replisome with RNA polymerase. To test this notion further, we determined the Spearman correlation rank matrixes for the gene excess for the predicted leading strand in the different codon positions, as well as for non-coding DNA (**Fig. 2b**, data was collected from the SkewDB). For the bacterial and archaeal replicons analyzed, this analysis revealed a strong correlation for an excess of G in the first codon position with correlation coefficients 0.85 and 0.92 for bacteria and archaea, respectively. The opposite trend was observed for the second codon position in both positions. These observations make sense in terms of the current understanding of genetic code and amino acid constraints^18^. Indeed, guanosine at the first codon position is a preferred nucleotide, whereas U/T and A at the second position are preferentially used to encode hydrophobic and hydrophilic amino acids, respectively. Therefore, biased gene density on the leading and lagging DNA strands contributes to the bacterial and archaeal GC skews. On the other hand, in bacteria, the contribution of G excess to GC-skew at the third codon position and non-coding DNA reflecting mutational biases are less strong. Very different results were obtained for archaea that demonstrate anti-correlation for the excess of G at the third codon position and the non-coding leading strand. Both of these observations are expected to decrease the G content on the leading strand and, consequently, the amplitude of the GC skew in archaeal species. Consequently, the relative contribution of the strand-biased gene density to archaeal GC skew must be relatively more important than in bacteria. We note that the relatively short size of Okazaki fragments in archaea^19,3^ may limit strand-specific cytosine deamination at the replication fork.

**Figure 2.**
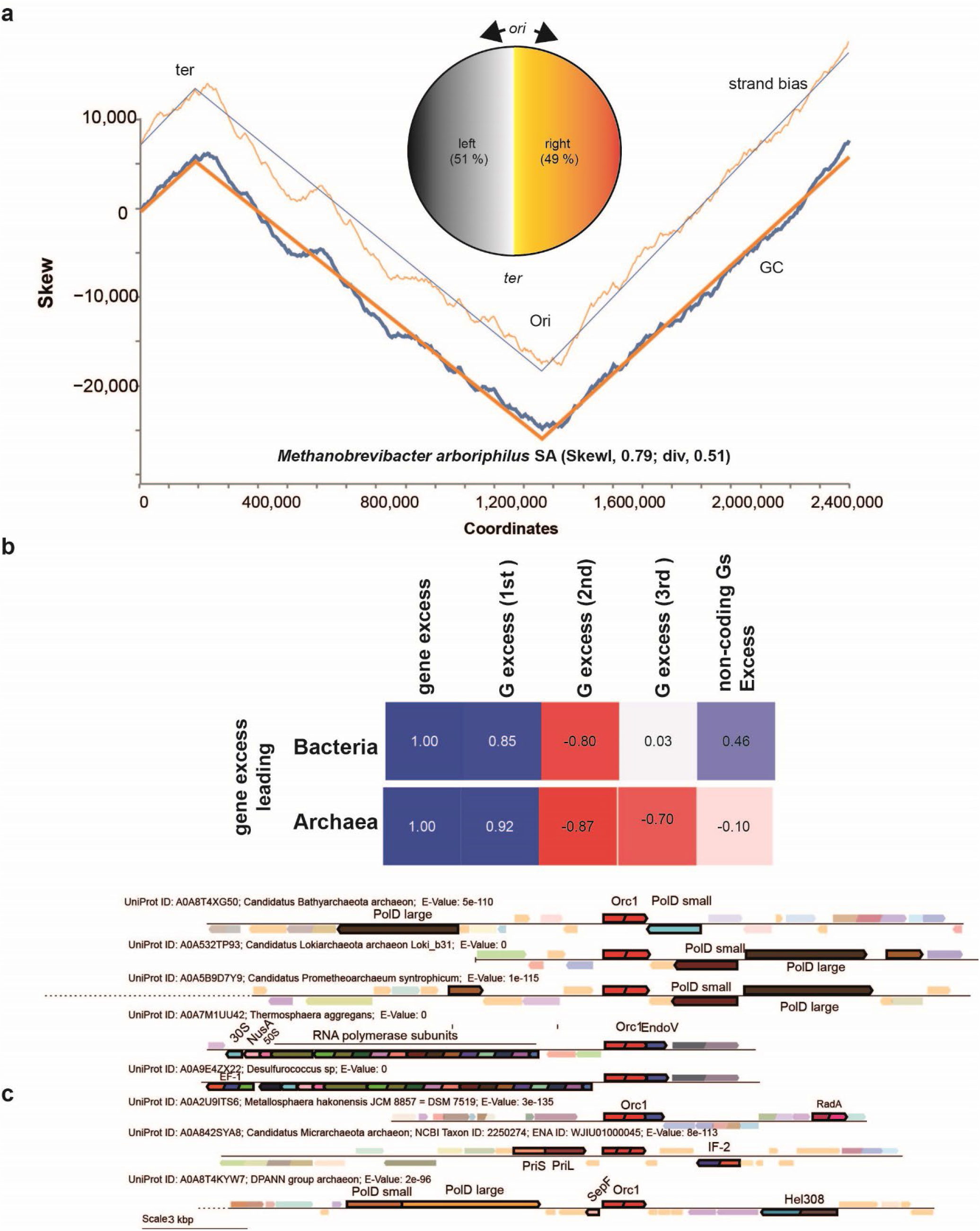
Genome-wide characteristics of the selected archaeal chromosomes. ***a)*** Example of the “ideal” archaeal chromosome with a quasi-perfect bacteria-like GC and transcriptional strand bias skews. ***b)*** Spearman correlation ecoefficiencies between the gene excess on the predicted leading strand of archaeal chromosomes with G excess in the indicated codon positions or non-coding DNA. ***c)*** Examples of statistically significant genome neighborhood associations with archaeal *orc1* encoding the replication initiator protein. Associations and their statistical significance was determined using EFI-Genome Neighborhood Tool that places protein families into a genomic context^21^. For details, see text.

Our results (**Fig. 2**) suggest that transcription and/or translation, and not replication, shape the archaeal genome nucleotide composition. This observation raises a possibility that the genetic and cellular organization of archaeal cells is a key aspect of the transfer of genetic information, similar to what has been suggested for bacteria^20^. To provide additional support for this poorly understood aspect of the archaeal chromosomes, we investigated the genomic contexts of the archaeal *orc1* genes. **Fig. 2a** provides examples of how *orc1* associates with DNA replication (DNA polymerase and primase subunits), DNA repair (NucS/EndoMS^12^, EndoV, Hel308), RadA recombinase, RNA polymerase subunits transcription, translation factors (50S, 30S, IF-2), and cell division (SepF) genes. Interestingly, these highly significant associations are found in a wide range of archaeal phylogenetic groups, including lokiarcheota, suggesting that the observed gene clustering is of general interest. Our protein-protein network analyses (**Fig. 3**) also revealed robust and significant interactions of *Pyrococcus abyssi* DNA replication and repair proteins Orc1, PCNA and the post-replicative mismatch repair endonuclease NucS/EndoMS with recombination proteins, the histones, and RNA polymerase subunits. Different DNA repair systems frequently have overlapping functions and must act in a coordinated and tightly regulated manner with other types of cell machinery. Therefore, understanding the evolution, function, and regulation of the detected interactions (**Fig. 3**) may not only increase our understanding of archaeal genome composition but also provide insight into how archaeal DNA repair systems could be used to increase the efficiency and specificity of modern genome editing tools.

**Figure 3.**
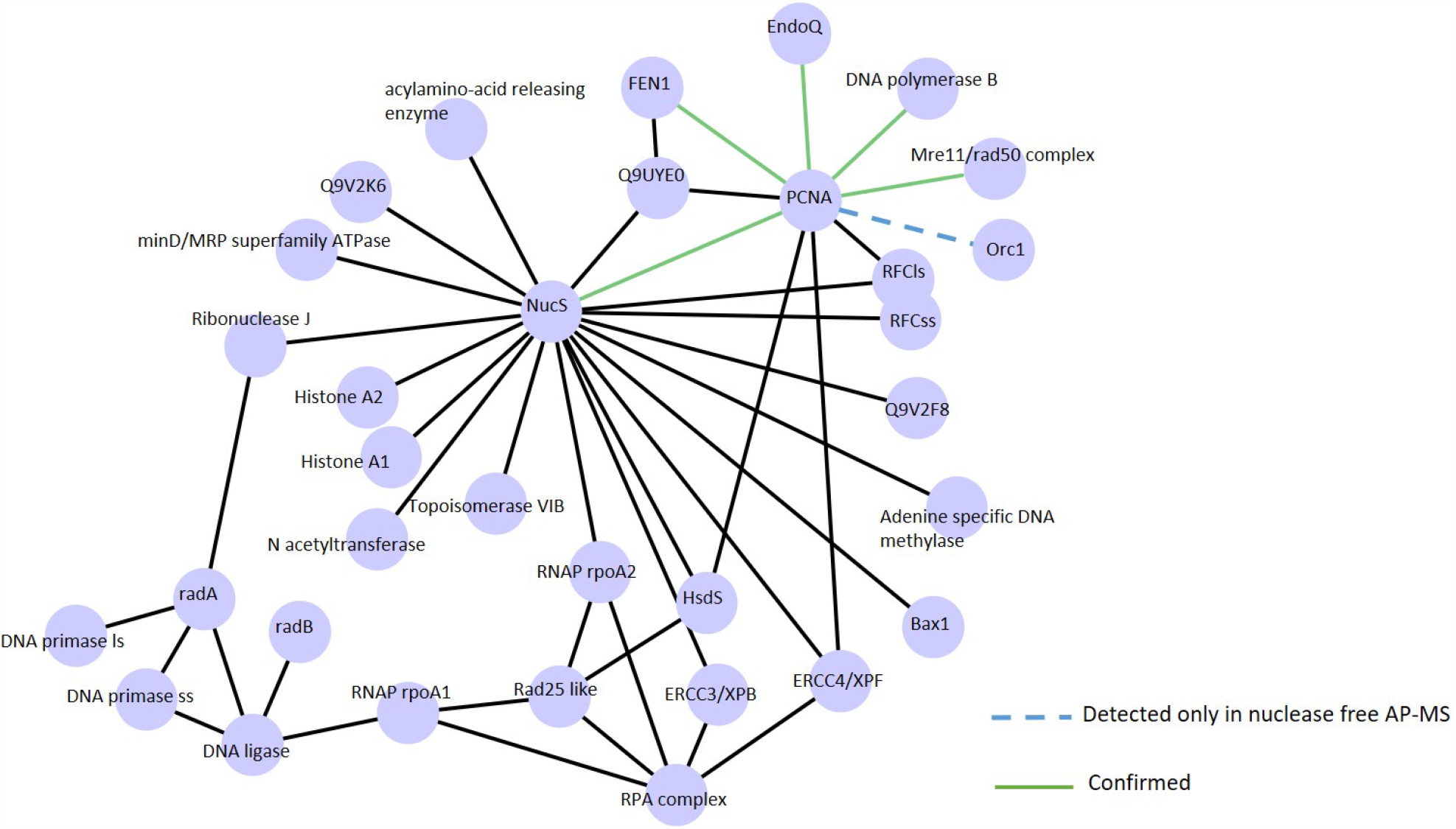
Systematic in vitro pulldown analyses of archaeal protein-protein interactions network analyses (for technical details, see^22^). This analysis revealed physical associations of the *Pyrococcus abyssi* DNA replication and repair proteins Orc1, PCNA and the post-replicative mismatch repair endonuclease NucS/EndoMS with recombination proteins, the histones, and RNA polymerase subunits.

In conclusion, our quantitative genome-wide analyses of the universally conserved GC-skew phenomenon revealed novel insight into evolutionary forces and molecular mechanisms that shape the nucleotide composition of archaeal genomes.

